# Genome-wide recombination map construction from single individuals using linked-read sequencing

**DOI:** 10.1101/489989

**Authors:** Andreea Dréau, Vrinda Venu, Ludmila Gaspar, Felicity C. Jones

**Affiliations:** Friedrich Miescher Laboratory of the Max Planck Society, Max-Planck-Ring 9, 72076 Tübingen, Germany

**Keywords:** recombination, crossover, meiosis, recombination hotspot, linked read sequencing, 10X genomics, mouse, stickleback.

## Abstract

Meiotic recombination is a major molecular mechanism generating genomic diversity. Recombination rates vary across the genome, often involving localized crossover “hotspots” and “coldspots”. Studying the molecular basis and mechanism underlying this variation within and among individuals has been challenging due to the high cost and effort required to construct individualized genome-wide maps of recombination crossovers. In this study we introduce a new method to detect recombination crossovers across the genome from sperm DNA using Illumina sequencing of linked-read libraries produced using 10X Genomics technology. We leverage the long range information provided by the linked short reads to phase and assign haplotype states to each DNA molecule. When applied to DNA from gametes of a diploid organism, the majority of linked-read molecules can be used to faithfully reconstruct an individual’s two haplotypes present at each location in the genome. A valuable rare fraction of molecules that span meiotic crossovers between the two chromosome haplotypes can then be isolated from the broader population of nonrecombinant molecules. Our pipeline, called ReMIX, allows us to characterize the genomic location and intensity of meiotic crossovers in a single individual and faithfully detects previously described recombination hotspots discovered by studies using mapping panels in mice. With a median crossover resolution of the mouse and stickleback being 15kb and 23kb respectively, ReMIX provides a powerful, high-throughput, low-cost approach to quantify recombination variation across the genome opening up numerous opportunities to study recombination variation with high genomic resolution in multiple individuals. ReMIX source code is available at at https://github.com/adreau/ReMIX.

---

Recombination is an essential process during meiosis. Chromosome segregation often occurs through crossing-over, which involves reciprocal exchange among homologous chromosomes and plays an essential role in meiotic chromosome segregation in sexually reproducing organisms. By shuffling parental alleles to produce novel haplotypes it is also a key source of genetic diversity that has considerable implications for the genomic landscape of variation and the evolutionary process.

In most diploid organisms, recombination is functionally constrained by the necessity for at least one well-positioned recombination event per homologous chromosome pair (this ensures proper segregation during Meiosis I) (Fledel-Alon et al., 2009). Defective, excessive, or deficient recombination can cause inviable gametes and developmental abnormalities (Hassold and Hunt, 2001, Inoue and Lupski, 2002). For these reasons the number of crossovers and their genomic locations are thought to be tightly regulated and highly constrained (Bernstein et al., 1994).

However, despite this core functional constraint, recent studies have revealed remarkable variation in recombination at multiple different scales (between and along chromosomes, among individuals, sexes, populations and species/taxa) (Comeron et al., 2012, Coop et al., 2008, Koehler et al., 2002, Kong et al., 2010, Nachman and Payseur, 2012, Paigen et al., 2008, Ptak et al., 2005). Crossovers are not uniformly distributed across the genome and the frequency (*recombination rate*), can vary by orders of magnitude and involve genomic *hotspots* and *coldspots*. For example, a well-studied recombination hotspot (*hlx1*) on mouse chromosome 1 has a remarkably high recombination rate of 2.63cM within a narrow 2.8kb interval in F1 hybrid male mouse (C57BL/6JxCAST/EiJ), yet is relatively colder in females of the same background and among other strains since the activity depends on the allele of recombination modifier protein *Prdm9*. Conversely recombination coldspots with a lack of crossovers in genomic regions as large as 6Mb have also been reported (Coop et al., 2008, Dumont et al., 2011, Lloyd et al., 2018, Paigen et al., 2008).

Part of this extensive variation may stem from the impact of recombination on individual fitness and rates of adaptation in natural populations - in addition to its fundamental role in meiosis, recombination impacts the inheritance of linked alleles, and may be subject to different selection pressures in different populations and different taxa. Depending on the evolutionary context, recombination may be beneficial if it breaks down linkage between deleterious and beneficial alleles (known as the Hill-Robertson Effect (Felsenstein, 1974, Hill and Robertson, 1966)), or deleterious if it breaks linkage between two adaptive alleles (Kirkpatrick and Barton, 2006). This variation in recombination has the potential to substantially impact genomic variation, individual fitness, and adaptation in natural populations, yet is poorly characterized and understood.

With the knowledge that number and genomic location of recombination can influence the segregation of traits, fitness of an organism, and adaptation in natural populations, there is increasing interest in the fields of medicine, agriculture and evolutionary genomics in the empirical quantification of fine scale variation in recombination among individuals, populations and species. Despite diverse approaches (linkage-maps, high density genotyping of pedigrees and individual sperm typing/sequencing), empirically quantifying recombination variation within and among individuals remains a challenge due to the expense and data intensity required to build numerous individualized genome-wide maps of recombination rate (Broman et al., 1998, Carrington and Cullen, 2004, Dumont et al., 2011, Kauppi et al., 2004, Kong et al., 2002, Li et al., 1988, Paigen et al., 2008, Shifman et al., 2006, Smeds et al., 2016, Wang et al., 2012). Other less data intensive approaches, such as comparisons of recombination among taxa using statistical estimates of recombination from population genetic (polymorphism) data, provide population and sex-averaged historical estimates of recombination rate and can be confounded by differences in the demographic history of the taxa and differences in the effective population size of the local genomic regions being compared. Further, these estimates make genetic dissection of molecular mechanisms underlying recombination variation difficult. In this study we address these challenges by introducing a new and powerful low-cost method that quantifies empirical recombination events across the genome of a single individual using linked-read sequencing of gametes.

Linked-read libraries are generated from long (high molecular weight) DNA molecules using a 10X Genomics Chromium controller. Numerous short reads are produced from DNA molecules trapped inside nanoliter sized droplets. Using their droplet-specific barcode these short reads can be computationally reconstructed into single molecules after Illumina sequencing. This low-cost long-range information can be used to solve the problem of haplotype determination. Our pipeline called ReMIX mines the long-range information in linked-read data to identify recombination crossovers across the genome. ReMIX makes use of some parts of the 10x Genomics pipeline, Long Ranger (Zheng et al., 2016), but deviates from it in a number of important ways. Long Ranger aligns reads to a reference sequence, calls and haplotype phases SNPs, reconstructs molecules, and identifies indels and large scale structural variants. It makes use of molecules that have a high probability of assignment to only one haplotype phase. Molecules that contain reads of mixed haplotype assignment (some reads assigned to one haplotype while others are assigned to the alternate haplotype), are considered to be errors and are discarded. However, when sequencing linked-read libraries from gamete DNA these haplotype switching molecules can also represent a valuable fraction of molecules spanning meiotic recombination crossovers. ReMIX identifies these valuable molecules and is the first method to enable reconstruction of individualized genomic recombination landscapes using linked-reads.

The linked-read information is exploited by ReMIX during three steps: identification of high-quality heterozygous variants, reconstruction of molecules, and the haplotype phasing of each molecule. The molecules identified as recombinant are then used to build an individualized genomic map of recombination crossovers, enabling us to quantify recombination variation across the genome.

We demonstrate our method using gametic tissue from a hybrid mouse and a stickleback fish. Genetic maps, available for both organisms, allow us to evaluate the accuracy of ReMIX. To validate the precision of our pipeline, we also use samples from the somatic tissue of the tested individuals as a negative control, as well as simulated data to determine the sensitivity and specificity of our method in genomes with different levels of polymorphisms. Using data from only a single individual and without prior knowledge of polymorphic sites, ReMIX obtained results that follow the same pattern of the previously described recombination maps, but with considerably higher resolution of the detected crossovers and lower costs compared to previous methods.

## Results

The novel method and algorithm that we present in this study uses pooled gamete DNA as starting material and reliably identifies recombination landscape of an individual at the whole genome level. Here we report the complete pipeline and results obtained by applying our method to an individual C57BL/6NcrlxCAST/EiJ hybrid mouse and freshwater stickleback fish (*Gasterosteus aculeatus*). High molecular weight DNA (>40kb) was extracted from purified sperm cells and somatic tissue of both mouse and fish (spleen and kidney respectively). 10X Genomics linked-read genomic libraries were prepared on a Chromium controller and the resulting linked-read libraries were sequenced on an Illumina HiSeq3000 sequencer to approximately 170X coverage. Reads obtained from the sequencing were then processed through our ReMIX pipeline to identify recombinant molecules and quantify the genomic recombination landscape of each individual.

### Overview of the ReMIX algorithm

ReMIX requires linked-reads generated from haploid gamete DNA as input. As a product of meiotic division, a haploid gamete comprises of a single copy of each chromosome in the genome - products of reductional cell division that are recombinants of the diploid parental chromosomes. The majority of linked-read molecules originating from a given haploid gamete, will be assigned with high probability to one of the two parental haplotypes. A small fraction of molecules (those spanning recombination crossovers) will contain reads that switch between the haplotypes. (Fig. 1A).

**Fig. 1:**
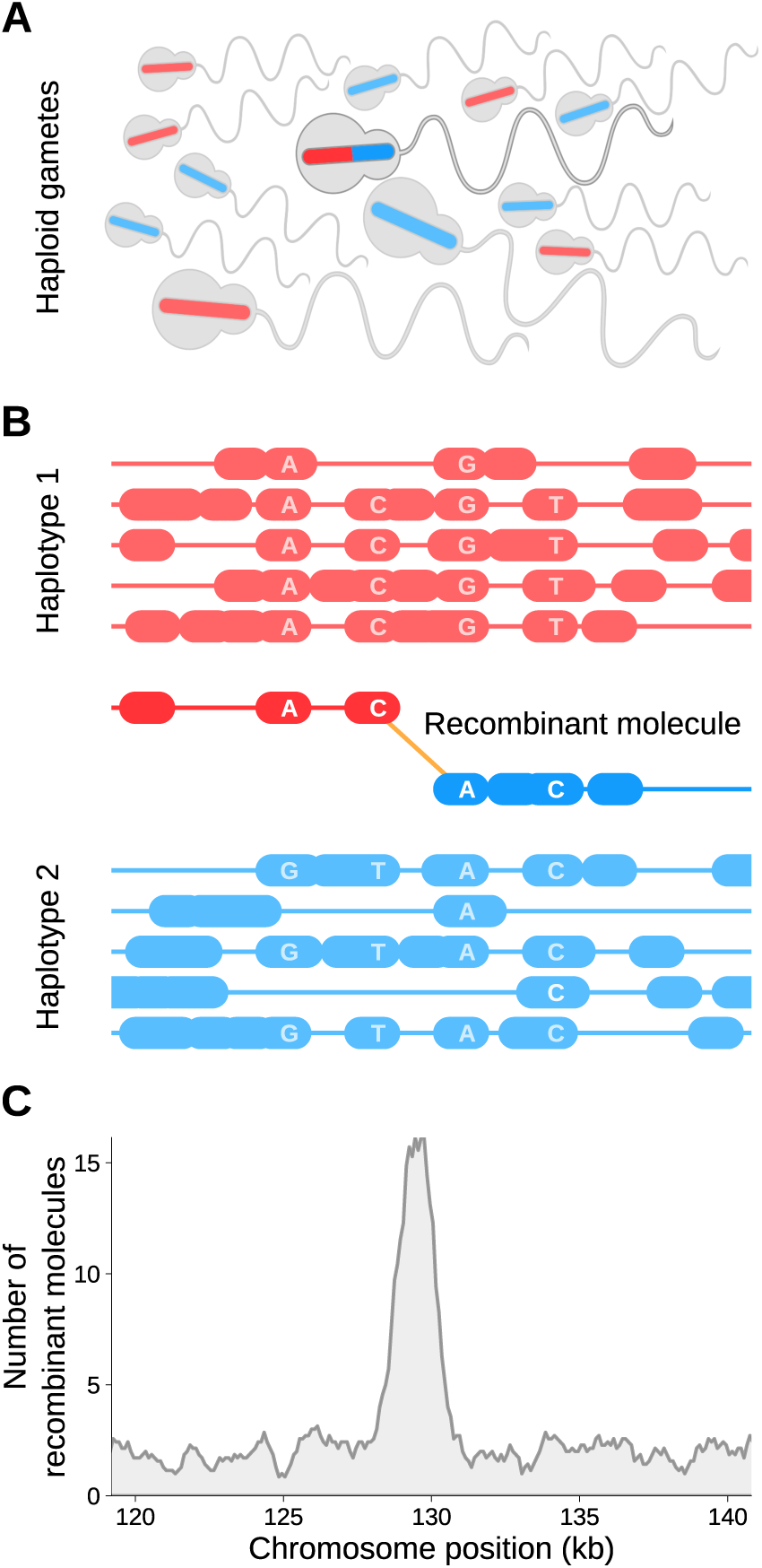
Construction of individualised genomic recombination maps using ReMIX. (A) DNA is isolated from a pool of sperm were each cell represents the haploid product of a single meiotic event. (B) ReMIX identifies high-quality heterozygous variants, reconstructs molecules then determines their haplotype phase. Three categories of molecules are identified: those belonging to haplotype 1 (red), haplotype 2 (blue), and recombinant molecules that switch from one haplotype to the other. Each contiguous line represents a molecule with the linked-reads marked by thick blocks. (C) Identified recombinant molecules are used to quantify the recombination rate across the genome.

The role of ReMIX is to identify the rare fraction of recombinant molecules as those that switch between haplo-types (Fig. 1B). For this, our pipeline aligns the linked-reads to a reference genome sequence in order to identify high-quality heterozygous variants and to reconstruct the original molecules.

Then the identified molecules are phased and separated into those that are entirely non-recombinant (haplotype 1 or 2 molecules), or alternatively, recombinant (haplotype switching) molecules. (full details in the Methods section).

A haplotype switching molecule may be generated from a true recombinant molecule or alternatively represent a false positive caused by bioinformatic errors such as sequencing error, incorrect read mapping, structural variation and barcode sharing among molecules from the same part of the genome. Our pipeline therefore incorporates several filtering steps to remove false-positive recombinant molecules. ReMIX initially filters the linked-reads based on the barcode sequence and the quality of the read. After variant calling the variants are filtered to remove polymorphisms showing allelic bias, and after molecule reconstruction, molecules with extreme high or low coverage are removed. Finally after the haplotype phasing of molecules, genomic regions that are not covered by a similar number of molecules for each haplotype are removed. These filters allow us to remove the regions that can introduce errors in the mapping or the phasing, such as copy number variation, small deletions, inversions, translocations etc. Finally, the ReMIX pipeline identifies molecules that have a high probability of containing a real crossover along with the genomic position of that crossover.

By considering the quality of each variant within a molecule, ReMIX is robust to small switches in haplotype state within a molecule when they are associated with low quality variant calls. This information is then used to build individualized genomic map of recombination crossovers (Fig. 1C).

### Identification of known hotspots in mouse

The genomic recombination landscape is well studied in various laboratory mouse strains, with one of the highest resolution sex-specific recombination maps constructed by (Paigen et al., 2008). Focusing on chromosome 1, the authors genotyped 6028 progeny produced from C57BL/6JxCAST/EiJ and CAST/EiJxC57BL/6J hybrids, mapped the locations of 5742 crossover events, and revealed the presence of a number of highly localized sex-specific recombination hotspots (Paigen et al., 2008). To evaluate the performance of our ReMIX pipeline, we analysed linked-read libraries produced from the sperm, and as a negative control, somatic tissue from the spleen, of a single C57BL/6NcrlxCAST/EiJ hybrid male. We compared ReMIX results with the high resolution recombination map from the 1479 C57BL/6JxCAST/EiJ male progeny (Paigen et al., 2008).

Whole genome sperm and somatic linked-read libraries were generated to sample a similar number of recombinant molecules on chromosome 1 as reported in (Paigen et al., 2008). We prepared 6 parallel reactions using the 10X Genomics Chromium controller each with ~1.2ng of DNA, approximately corresponding to a total of ~1700 haploid genomes. The final libraries were selected for an average of 600bp insert size and sequenced at 170x coverage with 2×150bp paired reads on an Illumina HiSeq3000. Both sets of linked-reads were analyzed using ReMIX and the latest version of the mouse reference genome, NCBI Build 38 (mm10) [GCF 000001635.20].

A crude estimate of the expected number of recombinant versus non-recombinant molecules can be made: for linked-read libraries made from a single gamete with an average molecule size of 60kb, sex-averaged map lengths of approximately 1630cM (genome-wide) and 96.55cM (chromosome 1) (Cox et al., 2009), and assembled genome size of 2.9Gb, we might expect to find recombinant molecules spanning crossovers at a frequency of 3.3 × 10^-4^ and 1.8 × 10^-5^ (16.3and 0.9 recombinant molecules in a genome-wide total of 48,333 molecules from a single gamete). In a pool of 1,700 gametes (equivalent to the number of gametes sequenced here), we expect to uncover 27,710 recombinant molecules across the genome, with roughly 1,641 of these located on chromosome 1.

After stringent filtering of the sperm sample ReMIX retained 1,210M reads and reconstructed 147,751,326 molecules with an average of 8 reads per molecule. A total of 30,508 (0.02%) molecules were identified as recombinant (genome-wide) and 2,369 of these were located on chromosome 1. Crossover positions of the recombinant molecules cluster into hotspots in a pattern closely mirroring the previously described male recombination map (Paigen et al., 2008) both in terms of position and intensity (Fig. 2A and Fig. S2).

**Fig. 2:**
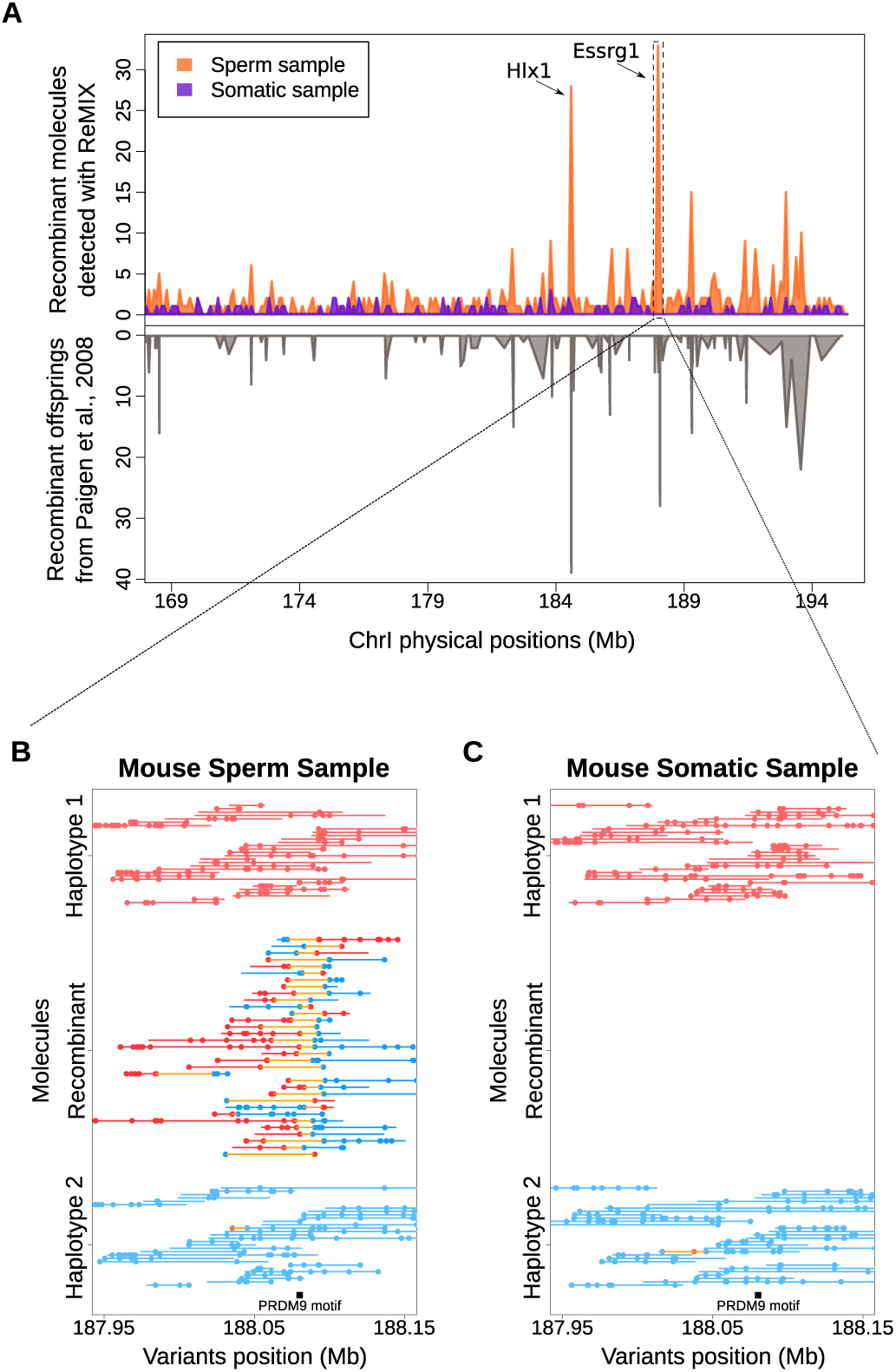
ReMIX correctly detects fine-scale recombination variation and hotspots on mouse chromosome 1. (A) The recombination rate on the south end of chromosome 1 (169-195.4Mb, mm10), determined by ReMIX corresponds well to the rate described in (Paigen et al., 2008). (B) The three types of molecules identified by ReMIX in the sperm sample in the region of a well-known recombination hotspot (Essrg-1, (Billings et al., 2013, Paigen et al., 2008)). Each line represents a single molecule and each dot a high quality heterozygous variant phased as haplotype 1 (red) or haplotype 2 (blue). For graphical reasons, we represented all the recombinant molecules detected by ReMIX but only 30 random classical molecules for each haplotype. (C) The corresponding region for somatic tissue lacks recombinant molecules. PRDM9 plays a role in initiating crossovers at the Essrg-1 hotspot and has a DNA binding motif (black bar) located near the midpoint of the detected recombinant molecules.

Accounting for false positives (See below), we see a number of windows that have significantly more crossovers than expected by chance (Fig. S10), suggesting the presence of hotspots in the mouse genome. In contrast, recombinant molecules detected in the somatic sample are less frequent, have a dispersed distribution and likely reflect false positives (discussed further below) from sequencing and/or bioinformatic errors (eg barcode collision) or rare mitotic recombination events. At the well known recombination hotspot region Essrg-1 reported by (Paigen et al., 2008) and (Billings et al., 2013) chr1:188,078,656-188,081,229 mm10 ReMIX identified 33 recombinant molecules in the sperm sample (Fig. 2B), while no recombinant molecules were identified in the corresponding genomic region in the somatic sample (Fig. 2C). Compared with previous studies involving more than 1500 mouse offspring, our results indicate that ReMIX is a powerful method for reconstruction of the fine-scale recombination landscape using gametes from a single individual.

We used both the number of recombinant molecules detected in the somatic sample and simulations (described in detail in below) to obtain independent estimates of the false positive rate. Adjusting ReMIX results according to these estimations, our data suggest the total number of true crossovers along chromosome 1 in 1700 sperm to be 1540, giving an average of 90.59 crossovers per meiotic product.

This corresponds well to the sex averaged genetic map length of mouse chromosome 1 (90.9cM), but is 8.3 cM longer than the map length of hybrid C57BL/6JxCAST/EiJ and hybrid CAST/EiJxC57BL/6J males: 81 and 83.65 respectively, calculated from Table S1 of (Paigen et al., 2008). This slightly higher number of observed recombinant molecules than expected based on the hybrid male map may have a biological basis (eg inter-individual variation (Koehler et al., 2002), inter-strain variation C57Bl6/J vs C57Bl6/Ncrl (Fontaine and Davis, 2016, Koehler et al., 2002), and possible differences arising from quantification of recombination from viable offspring vs quantification of recombination from gametes) or alternatively stem from detection errors (eg false negatives in the Paigen study due to lack of markers in the telomeric regions).

Finally, it has previously been shown that the genomic recombination landscape in mouse is positively correlated with CpG island density (Han et al., 2008). Here, we also find that recombinant molecules recovered by ReMIX are significantly closer to CpG islands than expected by chance based on 1000 (Wilcox rank sum test, *p <* 2.5 × 10^-20^)

### Finescale recombination landscape in stickleback fish

We next evaluated the performance of ReMIX in threespine stickleback fish - an evolutionary genomics model organism with reasonably high quality genome assembly, for which the recombination landscape has been described. (Glazer et al., 2015, Roesti et al., 2013, Sardell et al., 2018). To match the mouse sample, we created gametic and somatic linked-read libraries each using 0.8ng of high molecular weight DNA (approximately equivalent to 1700 gametes) from sperm and kidney tissue of a freshwater Scottish stickleback strain (River Tyne).

The libraries were selected for a mean insert size of 600bp and sequenced at 170X coverage on an Illumina HiSeq3000 machine. Both sets of linked-reads were analyzed using ReMIX and the stickleback reference genome (BROAD S1 (Jones et al., 2012), split into assembled scaffolds). 178M reads were retained post filtering and reconstructed into 21M molecules of which 2,639 (0,01%) were identified as recombinant by ReMIX.

The stickleback recombination landscape recovered with ReMIX corresponds well to the the rate inferred from the previous low resolution genetic map (Roesti et al., 2013) (Fig. 3 and S7). Consistent with previous studies, ReMIX reveals recombination crossovers are enriched towards the distal ends of chromosomes and are significantly clustered compared to random expectations (*p <* 1 × 10^-20^). Similar to the mouse results, ReMIX recovered a number of recombination molecules in the stickleback somatic sample providing an indication of a modest false positive rate (Fig. S8). For most chromosomes the maximum number of these false positive somatic recombinant molecules in 50 kb windows is 2 and we note some heterogeneity in the false positive rates across chromosomes with elevated levels on chromosomes XIV, XIX, and XXI (as high as 4 molecules on chrXXI) which co-localise with scaffold ends and are likely scaffold assembly errors.

**Fig. 3:**
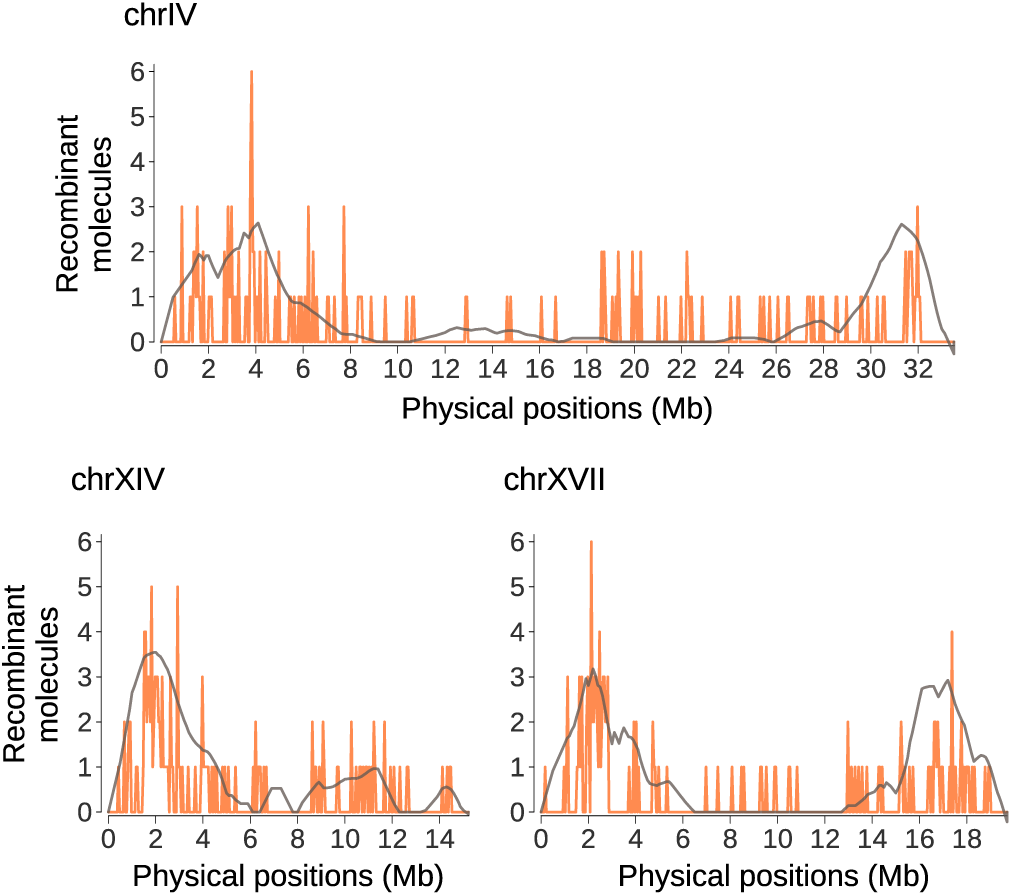
Recombination maps of three example autosomes in a male freshwater stickleback obtained using ReMIX analysis of linked-read data. The number of crossovers in 50kb windows identified by ReMIX (orange) is compared to recombination rate estimates obtained from a F2 lab cross (Roesti et al., 2013) of 140 males and 142 females individuals genotyped at 1872 markers (grey).

### ReMIX detects crossovers with high genomic resolution

A recombinant molecule is composed of two continuous sections: *s*_*a*_ phased to one haplotype and *s*_*b*_ phased to the opposite haplotype. The crossover may have occurred anywhere between the last informative variant of *s*_*a*_ and the first informative variant of *s*_*b*_. Thus we consider the resolution of a crossover as the physical distance between these two informative variants. By taking advantage of long range molecular data spanning high quality heterozygous variants segregating within a single individual, ReMIX directly identifies the recombinant molecules with high accuracy and crossover resolution (Figure 4). The achievable crossover resolution of our approach is limited primarily by the density of heterozygous sites within an individual (something that varies considerably across taxa), and secondarily by the sequencing coverage used to detect these informative sites. For example, based on whole genome sequencing data, we estimate hybrid C57BL/6NcrlxCAST/EiJ mouse and freshwater stickleback individuals used in this study will have a median distance of 44bp and 63bp between heterozygous sites respectively.

**Fig. 4:**
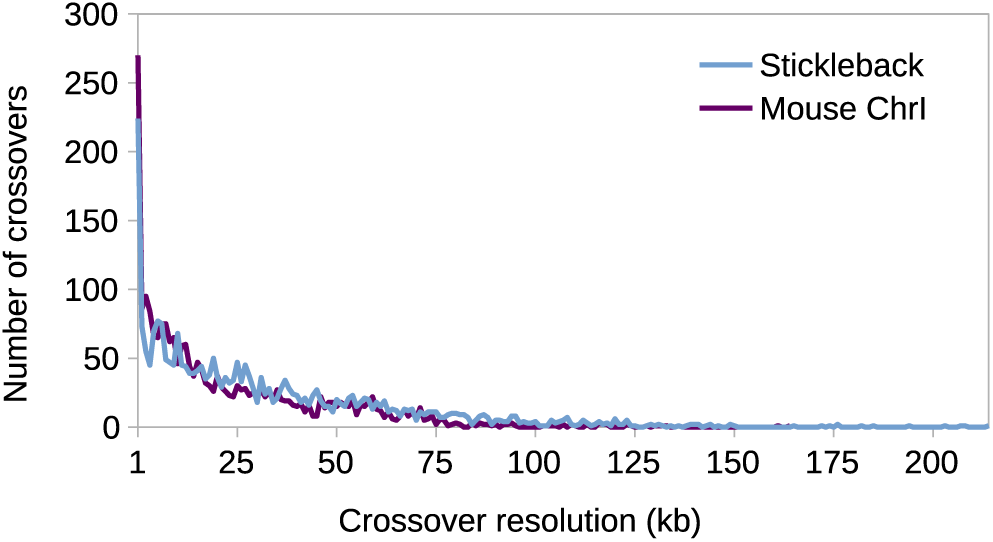
ReMIX detects recombination crossovers with high resolution in both mouse (purple) and stickleback (blue).

After stringent filtering of reads ReMIX achieved a mean of 8.3 and 8.5 reads per molecule, and a median crossover resolution of 14kb and 23kb for for mouse chromosome 1 and stickleback respectively. This is considerably higher than previous studies of mice (eg median resolution of 225 kb in (Paigen et al., 2008)) and close to the maximally achievable resolution based on the biological constraint of distance between heterozygous sites in these strains. The highest crossover resolution we achieved was 1 bp in both mouse and sticklebacks, while only 1.22% and 4% of the crossovers detected had low resolution of 100kb or more for mouse and stickleback respectively. We note that if desired, further improvements to crossover resolution up to the biological limit of distances between heterozygous sites could be achieved by increasing the depth of sequencing coverage (and consequently the number of reads per molecule).

### Analysis of accuracy on simulated data

Since fine scale recombination rate can vary considerably among individuals of the same species, comparisons of our ReMIX results above and previously published recombination provides only a qualitative assessment of the accuracy of our pipeline. To achieve a better indication of ReMIXs performance, we simulated several data sets using the linked-read simulator LRSIM (Luo et al., 2017). Starting from a reference sequence as an input, LRSIM can simulate diploid sequences with a user-specified number of heterozygous SNPs, indels and structural variants. Then the simulator extracts paired-end reads from each haplotype and then assigns the reads to molecules by attaching the specific 10x barcodes depending on a user-specified number of reads per molecule. In order to validate our method we generated linked-read sets containing both non-recombinant and recombinant molecules. To achieve this we first used LRSIM to create a set of linked-reads containing only non-recombinant molecules. Then we simulated crossovers between the two haplotypes (a switch of haplotype state) generated by LRSIM in the first run and we ran LRSIM on the recombinant haplotypes to obtain a second set of linked-reads containing both non-recombinant and recombinant molecules (those spanning the simulated crossovers). The resulting molecule sets were merged to simulate the mix of recombinant and non-recombinant molecules present in a pool of gametes. The sensitivity (or the true positive rate) is then computed as the proportion of the recombinant molecules correctly identified by ReMIX out of the total set of simulated recombinant molecules.

Let *m* be a recombinant molecule with two contiguous segments *s*_*a*_ and *s*_*b*_ phased to opposite haplotypes. ReMIX is able to detect *m* only if reads from both *s*_*a*_ and *s*_*b*_ are spanning heterozygous variants. Thus, the heterozygosity of the organism and the sequencing coverage are two parameters that influence the sensitivity of ReMIX to detect true positive recombinants. To evaluate the sensitivity we performed simulations with different heterozygosity levels and read density per molecule. The positions of heterozygous SNPs and reads were chosen randomly for each run. For each parameter configuration we ran the simulations ten times and averaged the sensitivity values. We show that ReMIX is highly sensitive (with more than 90% of recombinant molecules detected at moderate to high levels of heterozygosity and moderate to high sequencing depth (Figure 5).

**Fig. 5:**
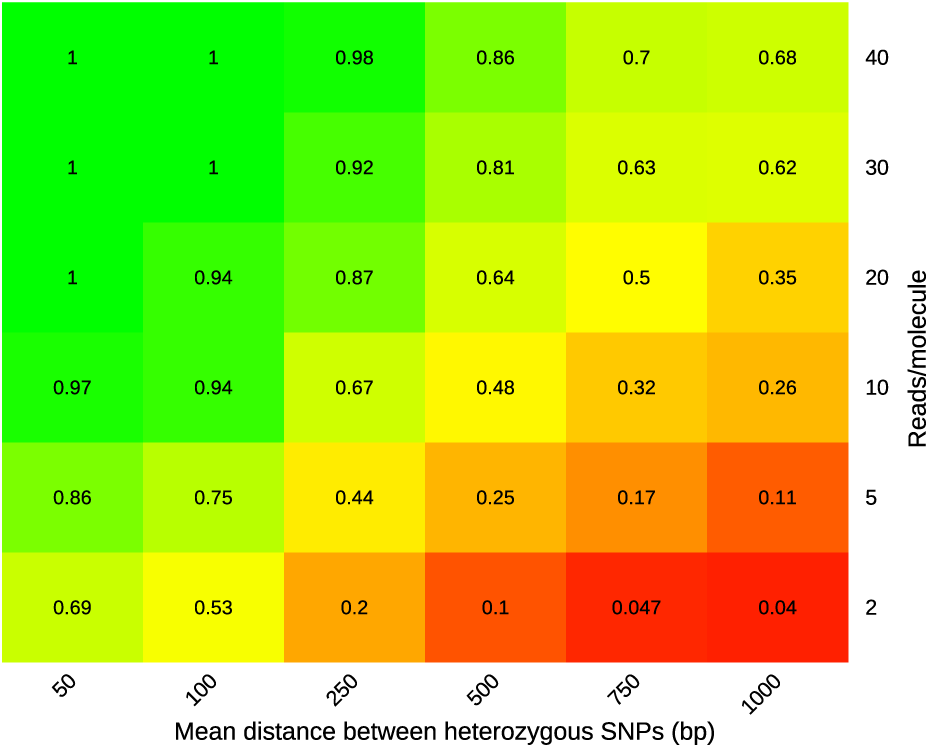
ReMIX detects recombinant molecules with high sensitivity at moderate to high heterozygosity levels and moderate to high sequencing depth.

The percentage of correctly reported molecules slowly decreases with the increase of the distance between the heterozygous variants. This is caused by the lower probability of reads spanning informative variants flanking recombination crossovers when an organism has a lower level of heterozygosity. However, ReMIX sensitivity can be easily increased in those cases by using a higher sequencing coverage.

Similar to other pipelines constructed for processing linked-reads (Weisenfeld et al., 2017, Zheng et al., 2016), the performance of ReMIX is dependent on the reaction conditions during the 10x Genomics linked-read library generation. One major consideration is the probability of two independent molecules from the same locus in the genome being assigned the same barcode (*barcode collision*). This depends on the amount of DNA in the reaction (which influences the number of molecules per droplet (GEM)), and the genome size of the organism in question. When preparing libraries from the same weight of DNA, small genomes will have a higher molecular copy number of each genomic locus, compared to large genomes. This leads to a higher probability of barcode collision of molecules from the same genomic locus due to a higher probability of them being trapped within the same GEM. In organisms with small genomes, using less DNA in the linked-read library preparation reaction can mitigate the occurrence of barcode collision.

If barcode collision occurs among alternate haplotypes, this has the potential to lead ReMIX to identify false positive recombinant molecules. Let *m*_1_ and *m*_2_ be two molecules of opposite haplotype state, from the same genomic region, that have the same barcode. The short reads are regrouped into molecules based on their barcode and a parameter specifying the maximal genomic distance separating two reads of the same molecule. Depending on the *m*_1_ and *m*_2_ read positions, the two molecules are detected as one molecule with two contiguous segments phased to opposite haplotypes. As a consequence, ReMIX reports the merged molecules as a recombinant molecule. We also note that, in some cases, barcode collision can facilitate the detection of recombinant molecules by ReMIX. When a recombinant molecule has strong disproportion between the numbers of phased variants mapping to each haplotype (example when only a few reads fall on one side of a crossover, while the majority fall on the other), collision with a second molecule representing the haplotype with low read coverage can help the identification of the recombinant molecule. Finally, identification of false positive recombinant molecules might also be caused by erroneous read mapping, structural variants, and reference genome assembly errors.

To address these two different causes of false positives, we used the complete mouse reference genome (mm10) to simulate a close to real case scenario for the numbers of molecules per GEM and the density and length of structural variants. Using the method described above, we simulated 7 molecules per GEM, the mean number of molecule per GEM that we obtained with our empirical mouse and stickleback data sets, and also 10 molecules per GEM, the maximum number reported by 10x Genomics. We then ran ReMIX on both sets and grouped the reported molecules in 100kb windows (Table 1). Under conditions matching our empirical datasets we estimated a low recombinant molecule false positive rate (with a large majority of the intervals not containing any false positive molecules (94.9%) and only 5% of intervals showing false positives). This increased to 10.37% of intervals when the number of molecules per GEM was simulated to be 10.

**Table 1:** Number of false positive (FP) molecules identified by ReMIX depending on the number of molecules per GEM. After spiting mouse genome in 100kb windows (total of 27,269), we reported the number of FP molecules identified by ReMIX in each window.

| No of FP per 100kb window | 0 | 1 | 2 | 3 | Total ratio of windows with FP |
| --- | --- | --- | --- | --- | --- |
| 7 mol/GEM | 25,889 | 1,335 | 44 | 1 | 5.06% |
| 10 mol/GEM | 24,440 | 2,678 | 145 | 6 | 10.37% |

The distribution of the intervals containing false positive molecules for mouse chromosome 1 for 7 and 10 molecules per GEM is shown in Figure S9 and S10 respectively. Similar levels of false positives were detected on the other mouse chromosomes. We note that due to the stringent filters of our pipeline (see Methods), structural variants were filtered, and did not have an impact on the false positive rate. Since the false positive molecules are uniformly distributed across the genome and do not cluster in specific regions, they do not interfere with the detection of regions with high recombination activity. And for organisms with low recombination rate the false positives detected by ReMIX can be decreased by lowering the amount of DNA used in the library preparation reaction, and the use of multiple independent reactions. This will decrease the mean number of molecules per GEM while maintaining the number of total recombinant molecules captured from the gametes.

## Discussion

Understanding the extent and molecular basis of recombination variation has been challenging due to the expense of creating individualized high-resolution genome-wide recombination maps. Here we present a cost and time effective method to build individualized recombination maps from pooled gamete DNA. This method makes use of linked-read sequencing technology developed by 10X Genomics to acquire long-range haplotype information from gametes of a single individual. Our specialized bioinformatics pipeline named ReMIX then faithfully identifies recombinant molecules from the linked-read data produced. Using these recombinant molecules, crossover locations are defined as genomic intervals based on the location of the last variant of the first haplotype and first variant of the second. We demonstrate the application of our method by building finescale recombination maps for a male mouse, an organism with well characterised recombination hotspots, and a less traditional model organism, a male threespine stickleback fish.

We validated our method through comparisons to previously reported recombination landscapes in mouse and sticklebacks. ((Paigen et al., 2008, Roesti et al., 2013)), and simulations to quantify sensitivity and specificity. Our approach faithfully identified known recombination hotspots on mouse chromosome 1 with high resolution (median of 14kb), and revealed enrichment in crossovers at the distal end of autosomes in the male mouse, and both ends of chromosomes in the male stickleback. Through simulations we show ReMIX has high sensitivity and that for organisms with low levels of heterozygosity this sensitivity can be increased by sequencing the linked-read library to higher coverage. In addition we used DNA extracted from somatic tissue as a control to test the specificity of our method. The use of a somatic control enabled the estimation of background noise in the data set that might be caused by bioinformatic error, reference genome assembly errors, copy number and structural variants, or rare mitotic recombination. Our results show that the true meiotic recombination signal stands out amidst the more dispersed noise from false positives, indicating ReMIX to be a reliable approach for constructing and studying variation in finescale recombination landscapes. Individualized genome-wide recombination maps that were previously constructed from extensive genotyping in thousands of offspring or whole genome sequencing of individual gametes (Wang et al., 2012) can now be produced with less time and effort by applying our novel method to pools of gametes from a single individual.

The whole genome recombination landscape we obtained for a male CASTxBL6 mouse (Figure S5) is in agreement with the reported observation that male recombination activity is concentrated at the distal end of the autosomes. We also detected previously reported mouse chromosome 1 hotspots (*Esrrg-1* and *Hlx1* (Fig. 2B and S3B) in our data set. Using a sliding window approach by counting number of haplotype switching molecules per 5kb interval, we find a 9kb interval at chr1:188,079,000-188,088,000 (mm10) region with highest recombination activity. This region spans the known Esrrg-1 hotspot. Crossovers were identified from 31 haplotype switching molecules out of 1736 total mapped molecules in that 9kb interval, suggesting a recombination rate of 1.78cM in 9kb. PRDM9, a protein with histone methyl transferase activity, plays an important role in recombination hotspots in many mammals including mice and humans. Consistent with previous studies showing that recombination in this region is mediated by PRDM9 ((Billings et al., 2013)), we find the PRDM9 motif specific to Esrrg-1 located within the 9kb hotspot (Fig. 2).

Mouse and stickleback recombination crossovers are not distributed randomly across the genome, but are rather significantly clustered and more proximal to CpG islands than expected by chance. In stickleback, the region with the highest recombination activity is located on chromosome IV at approximately 3.8Mb. Here, within a 7kb interval, we detected 6 recombinant molecules out of a total of 1366 mapped molecules. This corresponds to a recombination rate of 0.44cM, roughly one quarter the intensity of the mouse hotspot described above.

We have demonstrated our method here using DNA extracted from sperm in organisms with high quality genome assemblies. However, considering the ease of collecting pools of gametes, and the low amount of input DNA required (eg. 1ng for a genome size of 3GB genome, or less than 1ng for smaller genomes) we anticipate our method can be extended to a wide range of organisms. This opens up numerous research opportunities to explore variation in empirical finescale recombination maps in both model and non-model organisms including studying individuals sampled from the wild. For organisms whose genome assembly is of lower quality or lacking, a draft de novo assembly can be build based on the linked-reads set (Weisenfeld et al., 2017) required as an input for ReMIX and then used as a reference for ReMIX analysis of recombination. Further, with quick and reliable direct identification of recombination events in an individuals genome, our method also opens up the opportunity for forward genetic mapping of recombination modifiers in F2 panels - work that is ongoing in our lab.

## Methods

All animals used in this study were housed at approved animal facilities and handled according to Baden-Wrttemberg State approved protocols (Competent authority: Regierungsprsidium Tübingen, Germany; Permit and notice numbers 35/9185.82-5, 35/9185.46)

### Extraction of High Molecular Weight Genomic DNA

Stickleback genomic DNA was isolated from kidneys and sperm of a male wild-derived freshwater fish (River Tyne, Scotland). The sperm were collected via testes maceration in Hanks solution and purified to remove any potential contaminating diploid cells using a Nidacon PureSperm gradient following the manufacturers instructions. Purified sperm cells were resuspended in 1X PBS. Kidneys from the same male fish were dissected and rinsed in PBS prior to DNA extraction. High molecular weight gDNA was extracted from purified sperm cells and kidney using Qiagen Magattract HMW DNA extraction kit (Cat. No: 67563) following the protocol outlined in 10X Genomics Chromium Genome User Guide Rev B (10x Genomics, 2018). We followed the Genomic DNA extraction from cell suspension protocol for the sperm sample, and the Tissue DNA extraction protocol for the kidney sample.

Mouse genomic DNA was isolated from F1 hybrid (C57Bl6/NcrlxCAST/EiJ) male spleen and sperm cells. Sperm were collected from the cauda epididymis of a 7 week old F1 male hybrid mouse following (Ijiri et al., 2011). Extracted epididymides were finely chopped in 1xPBS (ThermoFisher, Cat. No:10010023). After settling for 3-5 minutes at room temperature, the supernatant containing viable sperm was purified by gradient centrifugation at 300 × g for 20min at room temperature (PureSperm 40/80; Nidacon International, Goteborg, Sweden). For somatic DNA control, excised spleen tissue was crushed between frosted glass microcope slides to make single cell suspension. Purified sperm and spleen cells were subsequently used for the isolation of high molecular weight (HMW) genomic DNA following (Wu et al., 1995).

The quality of extracted HMW DNA was checked by pulse field gel electrophoresis. All gametic and somatic samples showed a gradient of HMW DNA >50kb in size. This corresponds well to the described conditions for optimal performance of 10X Genomics linked-read library preparation (10x Genomics, 2018). No further size selection or processing was done on DNA samples.

### Sequencing library construction using the Chromium Genome Reagent Kit

We used a Chromium controller instrument (10X Genomics) to partition input DNA into nanoliter-sized droplets and pre-pare linked-read libraries following the manufacturers instructions (10X Genomics Chromium Controller User Manual) for input DNA quantification, dilution, GEM generation, and library preparation. For stickleback, we used ~0.8 ng of HMW genomic DNA as input (equivalent to 1700 haploid genomes). To achieve the equivalent number of haploid genomes for mouse (1700), we carried out 6 parallel reactions with 1.2ng input DNA for each of the sperm and somatic samples. In the Chromium Controller, input DNA was partitioned into ~1million droplets (GEMs), each containing reagents with a unique barcode (“Gemcode”). The droplets were recovered from the microfluidic chip and isothermally incubated (at 30 degree C) for ~3hrs to produce barcoded short reads, average size ~700bp, from each template DNA within each droplet. Following the isothermal incubation, the post GEM reads were recovered, then purified and size selected using Silane and Solid phase reverse immobilization (SPRI) beads. Illumina-compatible paired-end sequencing libraries were then prepared following the manufacturers instructions. The final library comprises reads with a standard Illumina P5 adapter, followed by a 16 bp 10X Genomics barcode at the start of read 1, the genomic DNA insert, and an 8bp sample index at the P7 adapter end. The final library was size selected to an average size of 600bp. Sequencing was conducted with an illumina HiSeq 3000 instrument with 2×150bp paired-end reads. Each library was sequenced to ~170X genome coverage.

### ReMIX pipeline for identification of recombinant molecules

ReMIX pipeline contains three main steps: identifying high-quality heterozygous variants, reconstructing molecules, and haplotype phasing each molecule to determine the recombinant molecules and the position of their crossovers (Fig. S1). We make use of the software provided by 10X Genomics for reference guided analysis of linked-read data (Long Ranger (Long Ranger, 2018)), but deviate from it in many places. After testing multiple equivalent tools for read filtering, alignment or variant calling, we have configured ReMIX with the combination of tools for which we obtained the best results using both simulated or real data.

### Identifying high-quality heterozygous variants

ReMIX’s detection of recombinant molecules is based on the estimation of the two haplotypes present in the diploid individual analysed. The accuracy of this estimation depends on the quality and frequency of heterozygous variants identified by our pipeline. Thus, in the first step of ReMIX (Figure S1) we remove the linked-reads containing sequencing errors in their genomic sequence, align the correct linked-reads on a reference genome, call the set of variants, and apply a hard filter on this set.

In step 1 of ReMIX (Figure S1 Step 1), the linked-reads are extracted from the Illumina’s sequencer base call files (*.bcl) using *Long Ranger mkfastq* (Long Ranger, 2018), and then filtered and trimmed with *Cutadapt* (Martin, 2011), *Trimmomatic* (Bolger et al., 2014) and *Long Ranger basic* (Long Ranger, 2018). The linked-reads with 16bp barcode sequences matching the barcode whitelist provided by 10X Genomics are aligned with *bwa mem* (Li and Durbin, 2009) to the reference genome. The duplicates are marked with *Picard tools* (Picard toolkit, 2018) and read alignment around indels is improved using *GATKs IndelRealigner* (McKenna et al., 2010). ReMIX identifies variants with *samtools mpileup* (Li, 2011) and applies a first variant filter using *bcftools* (Li et al., 2009) to extract high quality heterozygous variants with low allelic bias.

### Reconstructing the molecules

At the end of the first step of ReMIX the linked-reads are not yet organized into molecules. The purpose of the second step is to reconstruct the molecules so that the haplotype phasing algorithm can take advantage of the long-range information available.

The linked-reads generated from the same DNA molecule carry identical barcodes. However, since multiple molecules (eg 10) from diverse locations in the genome are typically trapped within the same GEM droplet and tagged with the same barcode, the molecules cannot be reconstructed based only on the barcodes of the linked-reads. From the quality control steps following HMW DNA extraction, it is possible to obtain an estimate of the expected average size of HMW DNA molecules in the reaction. Thus, we can link reads sharing an identical barcode into the same molecule if they aligned to the neighborhood of a genomic region with total molecule span similar to the expected average molecule size.

Still, this process does not always prevent linkage of reads from two or more independent molecules into a single reconstructed molecule when the original molecules share the same barcode and originate from the same genomic region. We refer to this case as *barcode collision*. For linked-read libraries constructed from organisms with large genome size using a low amount of input DNA in the library generation process, the probability of a single GEM droplet containing two HMW DNA molecules from the same genomic region is small, but non-zero. For example, the probability of barcode collision is approximately 3.2*x*10^-3^ for linked-read libraries prepared from 1ng of mouse DNA of 60kb average molecular weight, given a total mouse genome size of 3Gb (meaning approximately 3200 of 1M GEMs will contain more than one molecule from the same region of the genome). When the original molecules are generated from opposite haplo-types, the barcode collision cases can generate *recombinant-like* molecules that will be identified by ReMIX as false positive. To limit the number of false positives, we introduced the following parameters: the maximum molecule length, the maximum distance between two consecutive linked-reads grouped into the same molecule and the minimum and maximum number of expected linked-reads per molecule. The values of these parameters depend on the library construction and sequencing parameters.

For this second step of ReMIX we constructed a Long Ranger sub-pipeline called Long Ranger ReportMolecules (Figure S1 Step 2). This sub-pipeline is based on two parts of the Long Ranger Whole Genome Phasing and structural variant calling (SV Calling) pipeline (*Long Ranger wgs*) (Long Ranger, 2018): the computational reconstruction of the molecules, and the report of the molecule information in the INFO field of the variant call format (vcf) file. Long Ranger ReportMolecules incorporates a number of changes to the original Long Ranger pipeline including the parameters mentioned above: the maximum molecule length, the minimum and maximum number of expected linked-reads per molecule. The input of this sub-pipeline is the binary sequence alignment map (bam) file with high quality mapped reads including a tag with their respective barcodes, and the vcf file with the filtered heterozygous variants. Long Ranger ReportMolecules outputs a file that reports for each molecule: the genomic start and end position; the barcode; and the number of reads. This is accompanied by a modified vcf file that for each variant contains the reconstructed molecules spanning each of the alleles of this variant.

### Haplotype phasing the molecules

In the last step, ReMIX estimates the two haplotypes by phasing selected variants based on the molecule information previously obtained. Then, depending on the alleles spanned by the reads of a molecule, the molecule is considered as belonging to one of the two haplotypes or as being a recombinant molecule.

Structural variants such as deletions, duplications, copy number variations or translocations can cause errors in the read alignment, and thus variants can be incorrectly called in these regions. The false variants then interfere with the phasing process and introduce errors in the estimated haplotypes. Moreover, the structural variants can generate *barcode collision*-like cases. If misplaced reads and a real molecule share the same barcode and are aligned in the same genomic region, the algorithm used for reconstructing the molecules regroups the misplaced reads and the real molecule in a unique molecule. When the misplaced reads and the real molecule originate from opposite haplotypes, the reconstructed molecule appears as if it would span a crossover event as presented in Figure S11. ReMIX identifies these problematic regions by removing: variants that have a notable difference between the molecular or read coverage compared to the mean values for their chromosome; and variants for which the read coverage is uneven between the alleles.

The remaining variants are then phased with HBOP (Xie et al., 2012) based on the molecules computed during the second step. HBOP is a single individual phasing algorithm that can take into account reads belonging to a longer DNA fragment and therefore capitalizes on the long-range information of the molecule during phasing.

The two haplotypes constructed by HBOP are then used to phase each molecule. For each variant spanned by a molecule with at least one read, we consider the haplotype of the covered allele and the sequencing quality score at that position. Then, based on a score function implemented in Long Ranger wgs (Long Ranger, 2018), we compute for each molecule the probability of belonging to the two haplotypes or being a mix of the two. Contrary to Long Ranger wgs, we do not consider the molecules that contain reads spanning both alleles of a variant since this is behavior is likely to arise from a barcode collision. Once the probabilities are computed for each molecule, we filter again filter again to remove variants showing an allelic bias in the number of molecules phased to each allele. filter again to remove variants showing an allelic bias in the number of molecules phased to each allele Depending on the quality of the reference sequence used in the mapping process or on the copy number variation, some of the structural variants are still unidentified and can introduce errors in the process of determining the recombinant molecule. We then recompute the haplotype probabilities for each molecule.

From the set of molecules that have a high probability of belonging to a mixture of two haplotypes states, ReMIX considers as truly recombinant the molecules for which we can identify a clear crossover position: a minimum number of variants and a minimum ratio of variants phased to the same haplotype on each side of the crossover. We then output for each recombinant molecule the genomic start and end position; the crossover positions; the barcode; and the number of reads.

## Supporting information

## Software availability

ReMIX source code can be found at https://github.com/adreau/ReMIX.

## Acknowledgements

We would like to thank Frank Chan for input into experimental designs, useful discussions, and ideas, his contribution of mouse material. We also like to thank both Frank Chan and Marek Kucka for their insight and ongoing discussions related to further development of linked-read technology for recombination detection. We thank Elena Avdievich and Enni Harjunmaa for their suggestions and support on library preparation and data analysis. We are grateful to Ruth Ley for her contribution towards the 10X Genomics Chromium controller, to Andre Noll for his high performance computing support, and Christa Lanz for her assistance with high throughput sequencing. ReMIX uses part of the 10X Genomics Long Ranger pipeline and we are grateful to 10X Genomics for access to their open source code and their discussions in the initial stages of this project.

