## Supplementary material for "Genome-wide recombination map construction from single individuals using linked-read sequencing"

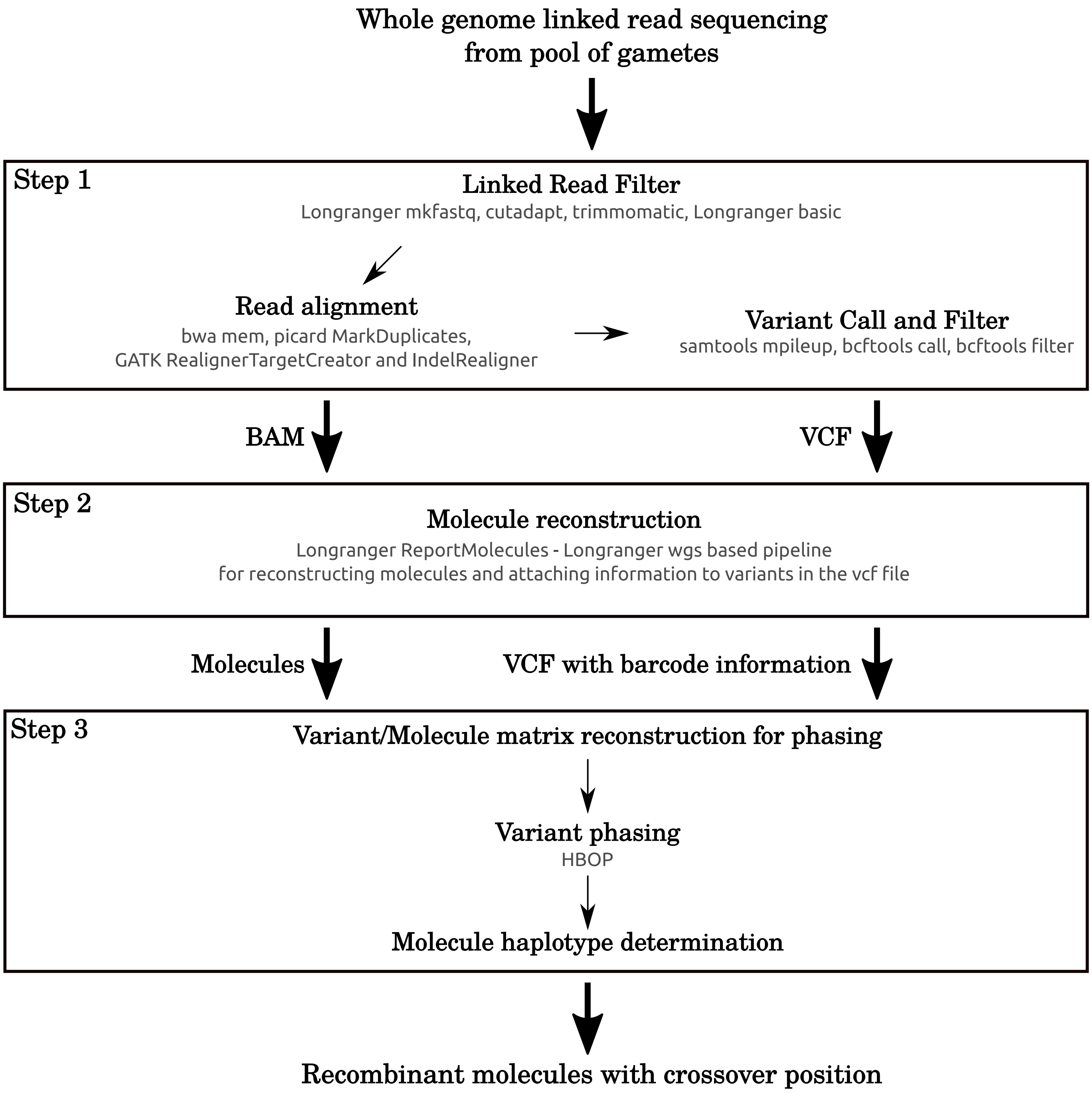


Fig. S1: ReMIX pipeline's main steps: identification of high-quality heterozygous variants, reconstruction of molecules, and haplotype phasing each molecule. In the input our pipeline requires Illumina's base call files from sequencing pool of gametes and outputs the identified recombinant molecules and the position of their crossovers.

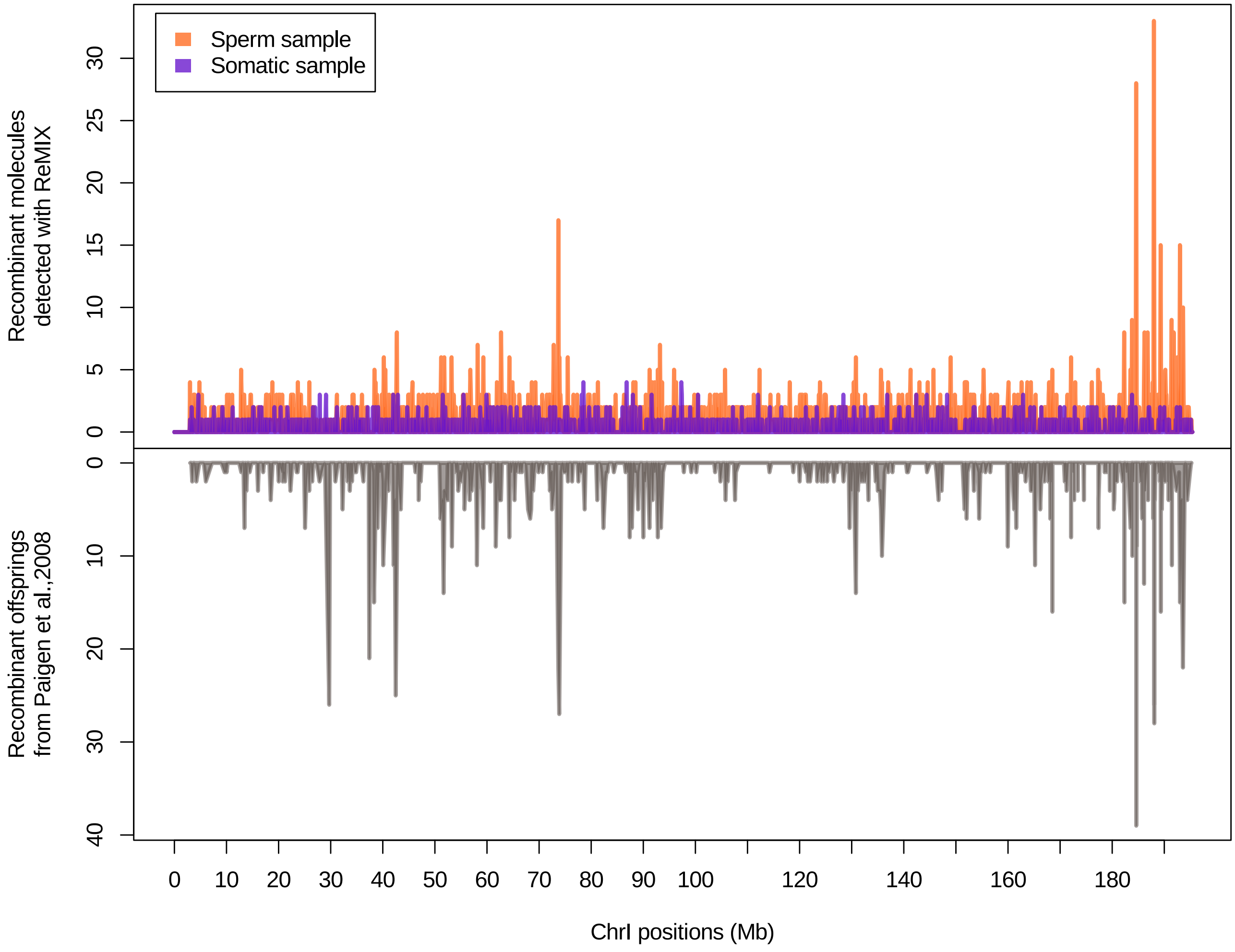


Fig. S2: ReMIX correctly detects fine-scale recombination variation and hotspots on mouse chromosome 1. The recombination rate determined by ReMIX corresponds well to the rate described in (Paigen et al., 2008). The dissimilarity observed in the northern end of the chromosome may be caused by potential sub-strain differences in recombination (Fontaine and Davis, 2016) (C57BL/6NcrlxCAST/EiJ used in our study and C57BL/6JxCAST/EiJ in (Paigen et al., 2008).)

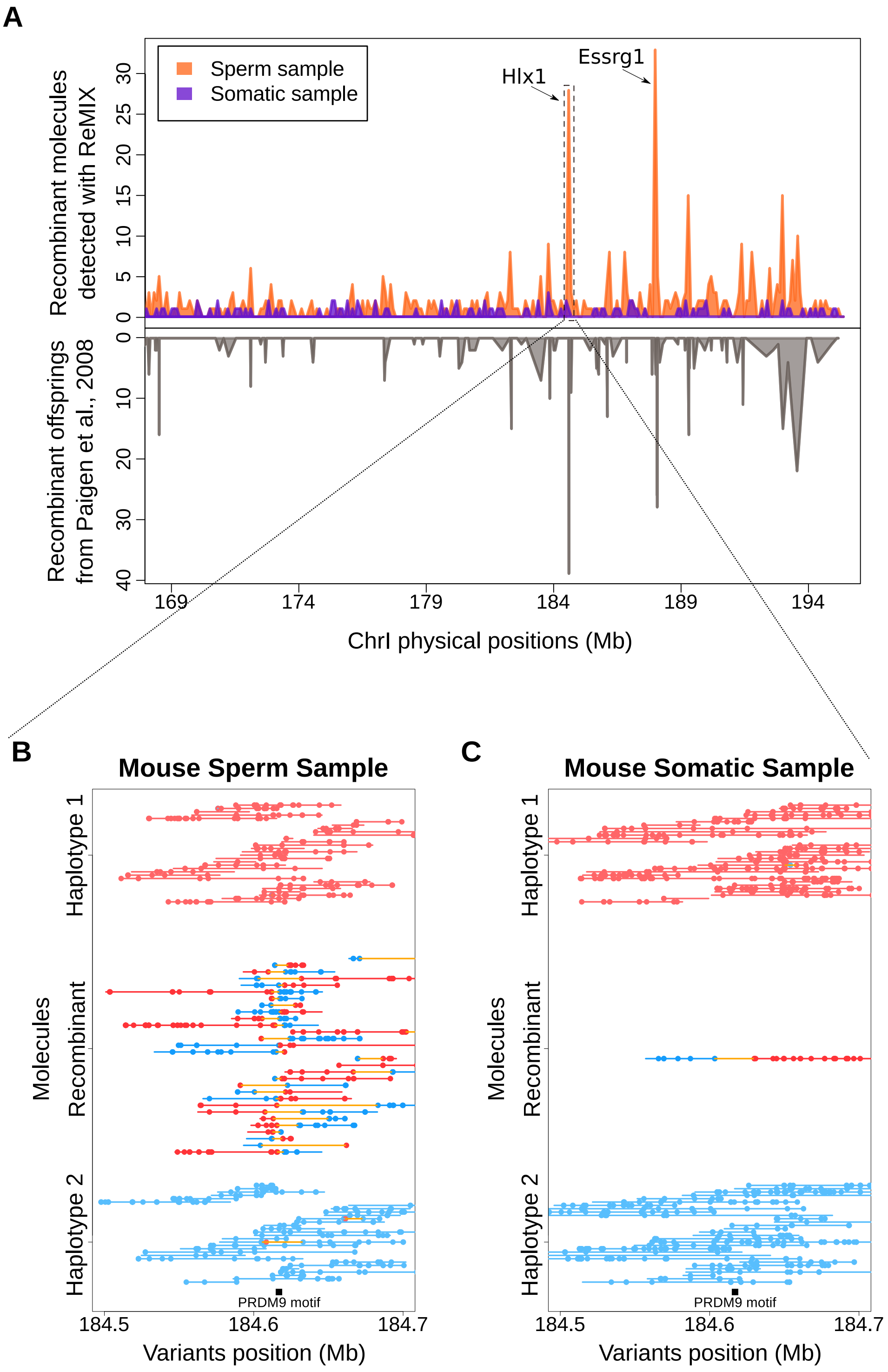


Fig. S3: ReMIX correctly detects fine-scale recombination variation and hotspots on mouse chromosome 1. (A) The recombination rate on the south end of chromosome 1 (169-195.4Mb, mm10), determined by ReMIX corresponds well to the rate described in (Paigen et al., 2008). (B) The three types of molecules identified by ReMIX in the sperm sample in the region of a well-known recombination hotspot (Hlx1, (Billings et al., 2013, Paigen et al., 2008)). PRDM9 plays a role in initiating crossovers at the Hlx1 hotspot and has a DNA binding motif (black bar) located near the midpoint of the detected recombinant molecules. (C) The corresponding region for somatic tissue in which ReMIX identified a recombinant molecule due to barcode collision.

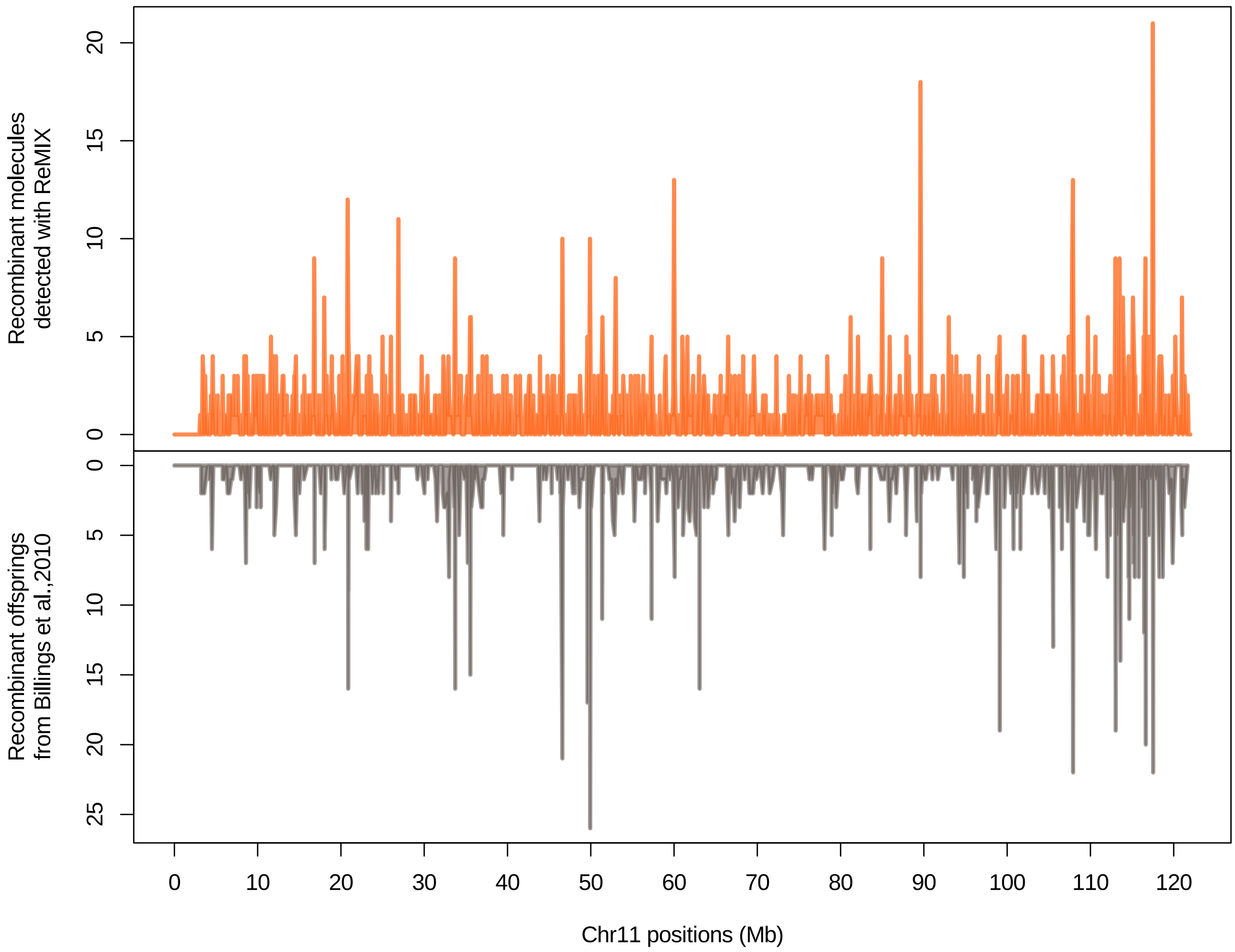


Fig. S4: ReMIX correctly detects fine-scale recombination variation and hotspots on mouse chromosome 11. The recombination rate determined by ReMIX corresponds well to the rate described in (Billings et al., 2010).

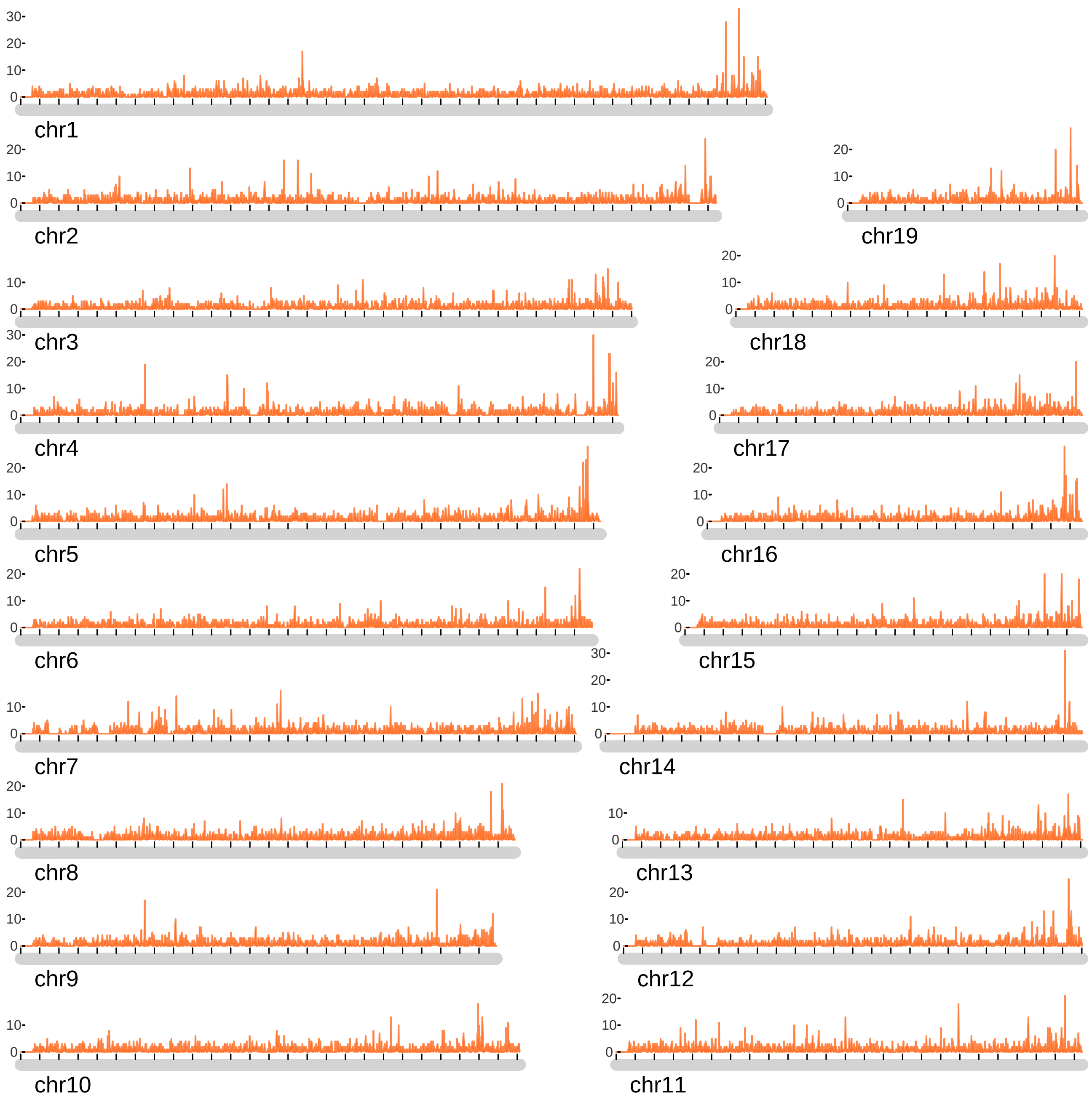


Fig. S5: ReMIX results on the mouse autosomes. Consistent with previous studies (Liu et al., 2014), ReMIX reveals recombination crossovers are enriched towards the distal ends of chromosomes in male germline.

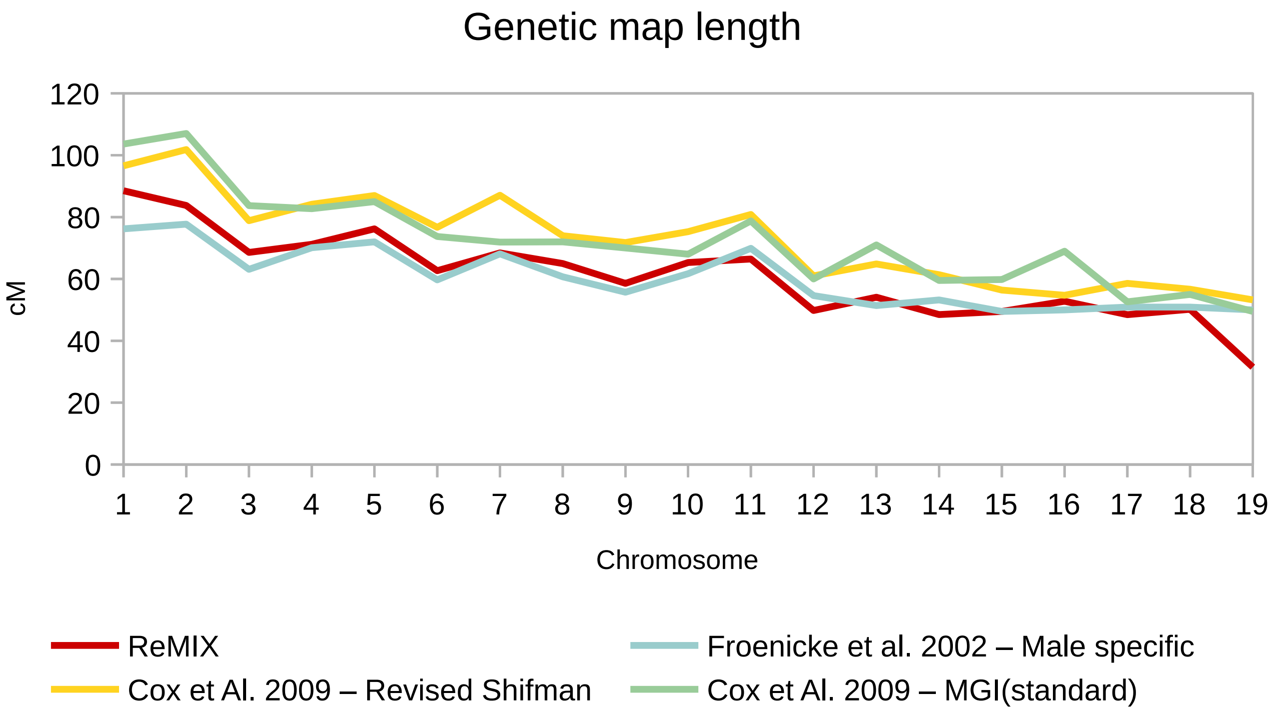


Fig. S6: Genetic map length comparison between previous studies analyzing various mouse strains and ReMIX results on the mouse genome.

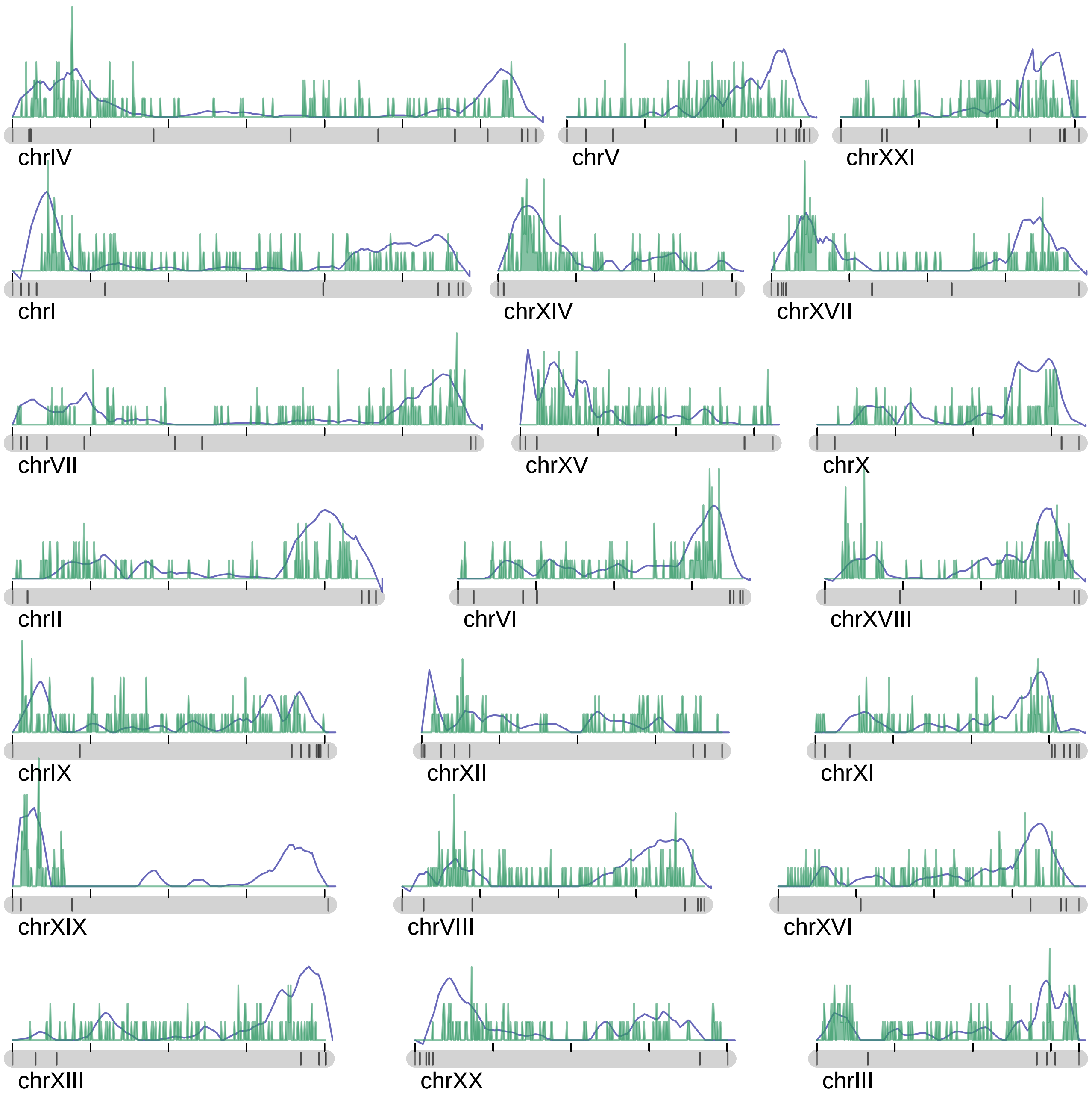


Fig. S7: Genome graph of recombination events in a male freshwater stickleback with underlying genetic map. The number of crossovers identified by our pipeline is plotted in 50 kb intervals (in green). The genetic map was previously constructed from F2 lab cross population of 282 male and female individuals and 1872 total markers (Roesti et al., 2013) (in blue).

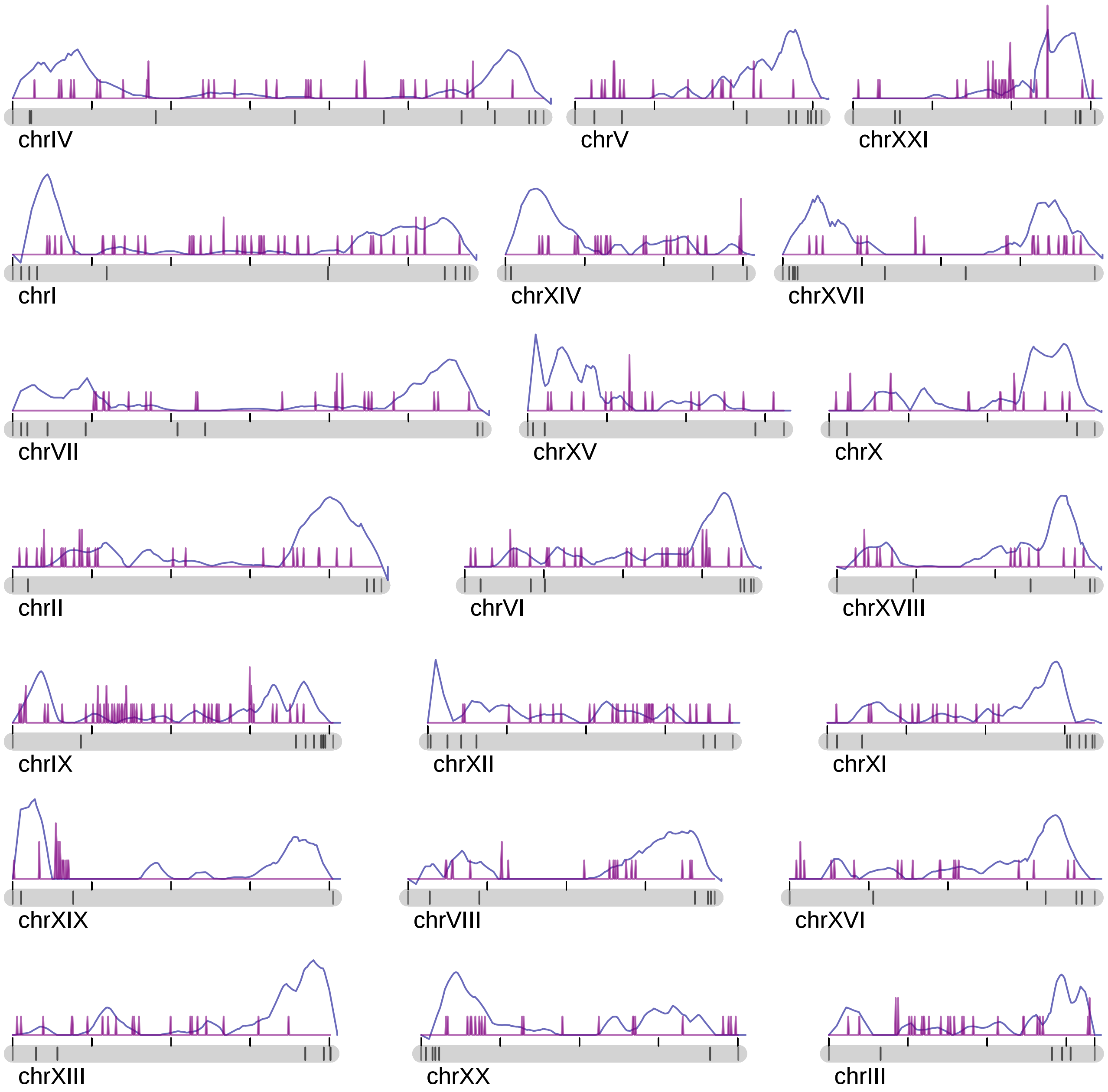


Fig. S8: Genome graph of recombination events in somatic tissue sample with underlying genetic map for negative control. The number of crossovers identified by our pipeline is plotted in 50 kb intervals (in purple). The genetic map was previously constructed from F2 lab cross population of 282 male and female individuals and 1872 total markers (Roesti et al., 2013) (in green). For most chromosomes the maximum number of these false positive somatic recombinant molecules in 50 kb windows is 2. As expected, the moderate false positive rate is evenly distributed and does not interfere with the hotspot detection. The false positive rates across chromosomes with elevated levels co-localise with scaffold ends (black lines on the gray bars) and are likely scaffold assembly errors .

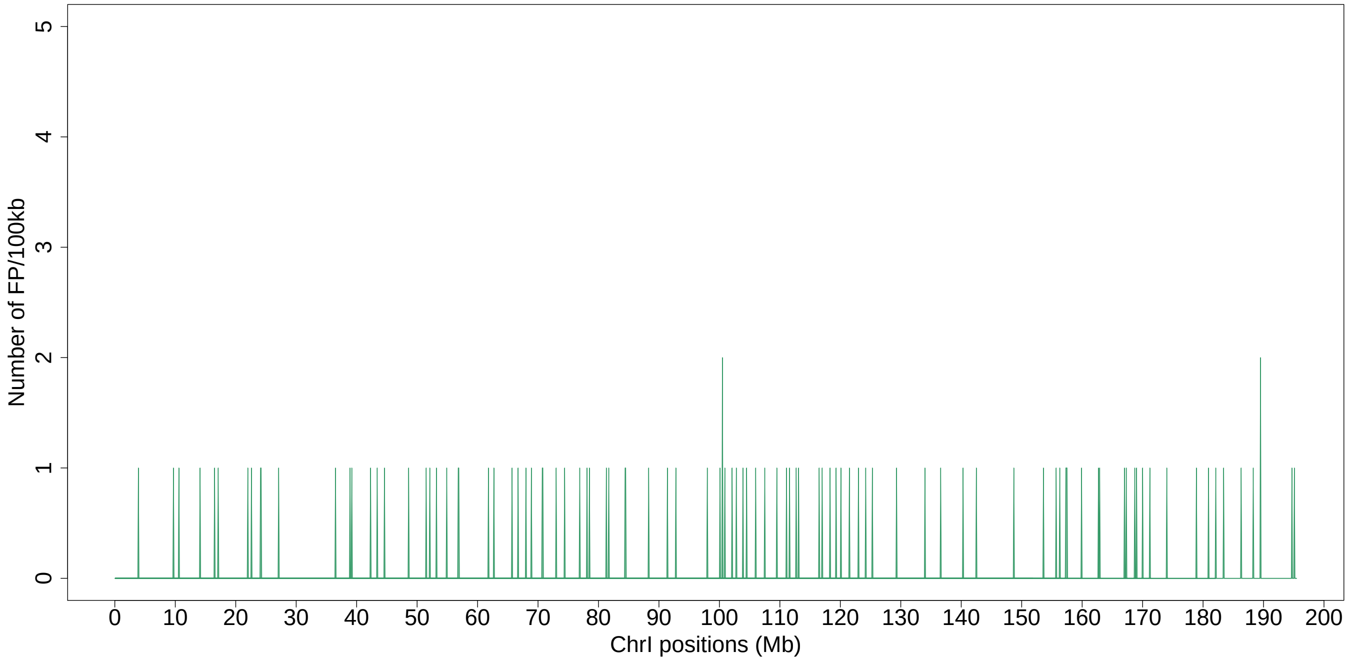


Fig. S9: Frequency of false positive molecules in 100kb windows in the case of simulating 7 molecules per GEM in the mouse chromosome 1.

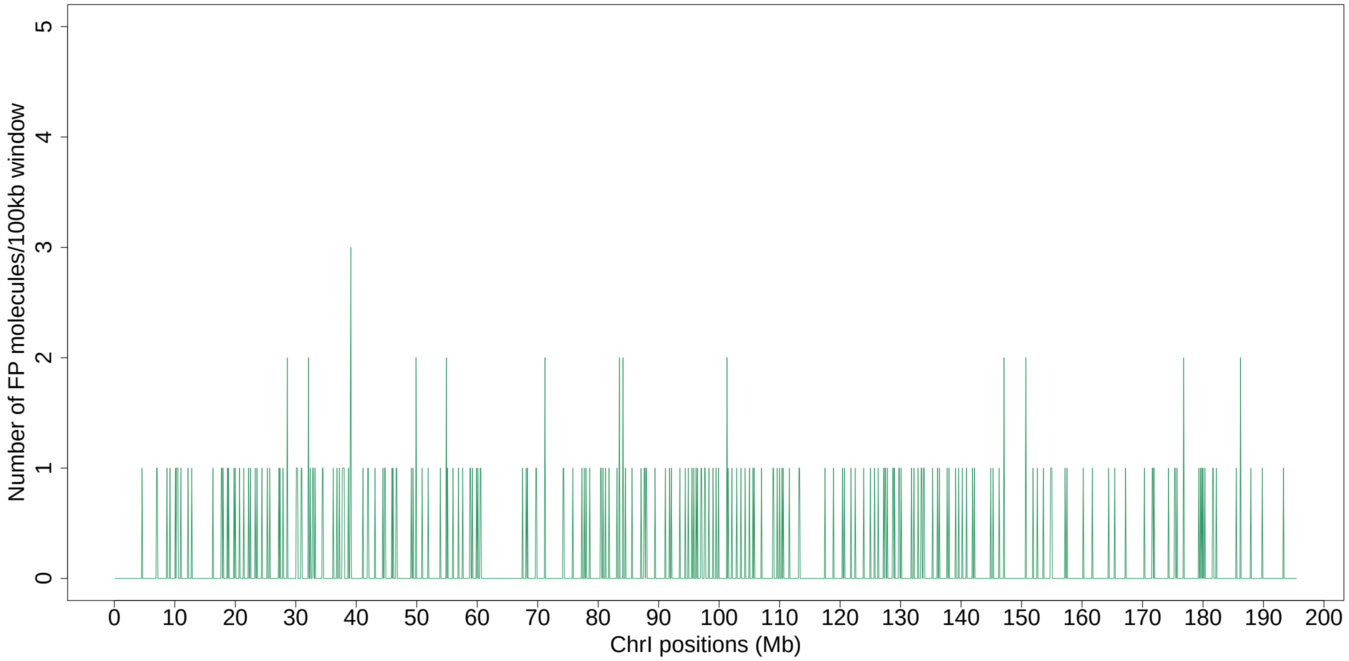


Fig. S10: Frequency of false positive molecules in 100kb windows in the case of simulating 10 molecules per GEM in the mouse chromosome 1.

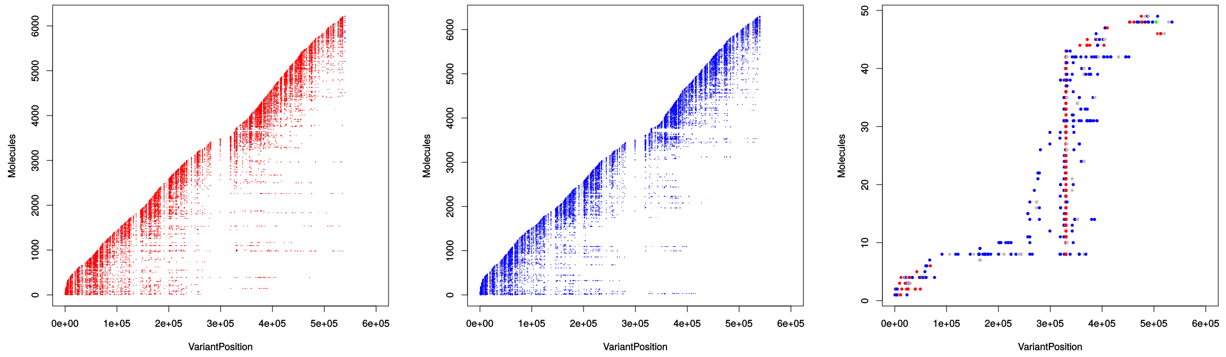


(a) Haplotype 1 molecules.

(b) Haplotype 2 molecules.

(c) Recombinant-like molecules.

Fig. S11: The errors in the read alignment and variant calling due to structural variants can generate recombinant-like molecules. These errors can cause incorrect variant phasing or barocode collision cases in the structural variants regions. When misplaced reads and a real molecule share the same barcode and are aligned in the same genomic region, the algorithm used for reconstructing the molecules regroups the misplaced reads and the real molecule in a unique molecule. In the case when the misplaced reads and the real molecule originate from opposite haplotypes or when variants are incorrectly phased, the reconstructed molecules appear as if it would span a crossover event and they pile-up wrongly suggests a hotspot region. ReMIX effectively identifies these problematic regions and removes them.
